# Spatially Organized IGF1-mTOR Signaling Controls Human Forebrain Progenitor Fate Through Coordinated Transcriptional and Translational Programs

**DOI:** 10.1101/2025.05.08.652851

**Authors:** Kadia Lisst, Da Huo, Stephen M. Eacker, Sonya Zhang, Alina Yang, Yuejia Huang, Ted M. Dawson, Valina L. Dawson, Jin-Chong Xu

## Abstract

The specification and maintenance of human forebrain neural progenitor cells (NPCs) depend on both intrinsic gene networks and spatially localized niche signals, but the interplay between these cues remains incompletely understood. Here, we identify a spatially organized, paracrine IGF1 signaling architecture that regulates human FOXG1⁺ NPCs through multilayered transcriptional and translational control. Using a pluripotent stem cell–derived forebrain model, we show that FOXG1⁺ NPCs express IGF1 receptors but lack endogenous IGF1, instead depending on neighboring epithelial-like domains that secrete IGF1. IGF1 promotes progenitor proliferation, clonal expansion, and vertical tissue growth by activating PI3K–AKT–mTOR and MEK–ERK pathways. Ribosome profiling and 5′UTR reporter assays reveal that mTOR signaling selectively enhances translation of neurodevelopmental and biosynthetic transcripts— including *GSX1*, a ventral fate determinant implicated in interneuron specification and autism. These findings uncover a human-specific regulatory mechanism in which spatially restricted IGF1–mTOR signaling integrates niche signals with translational output to support progenitor identity, biosynthetic capacity, and developmental resilience.

## Introduction

The expansion and maintenance of neural progenitor cells (NPCs) are essential for shaping the developing human forebrain, where regional patterning, progenitor pool size, and neurogenic timing must be precisely regulated^1^. Disruptions in these early processes are increasingly linked to neurodevelopmental disorders, including autism spectrum disorder^2^ and epilepsy^3, 4^. In TSC1-associated disease, for example, cortical tubers arise during prenatal development from dysregulated mTOR signaling in forebrain neural progenitor cells—prior to neuronal migration and cortical lamination^5–8^. In contrast, mTOR dysregulation in the postnatal or adult brain does not cause tuber formation, instead impacting plasticity, gliosis, or metabolic regulation^5, 9–13^.

These observations underscore the importance of temporally and spatially restricted signaling events in forebrain progenitor fate decisions.

While intrinsic transcriptional programs guide neural specification, the full range of extrinsic cues—and how they interface with intracellular signaling pathways in human forebrain NPCs— remains incompletely understood^14, 15^. Among the central regulators of cellular growth and developmental plasticity are the Insulin-like Growth Factor (IGF)^16^ and mechanistic Target of Rapamycin (mTOR) pathways^17^, which coordinate nutrient sensing, proliferation, and biosynthetic activity across diverse tissues^18^. Yet their specific roles and upstream regulation in regionally defined human forebrain progenitors have not been fully elucidated.

Given the inaccessibility of early human fetal brain tissue, pluripotent stem cell–derived models now offer an experimentally tractable platform to study these signaling pathways in developmentally relevant contexts^19^. While in vitro and in vivo studies in rodents have shown that IGF1 promotes neural proliferation and brain growth during early development^20, 21^, they lack human-specific forebrain patterning and do not address niche architecture or translational regulation in FOXG1⁺ progenitors. Our study addresses this gap by investigating how IGF1– mTOR signaling governs human forebrain NPC behavior within a spatially organized, paracrine niche.

Using a human pluripotent stem cell–derived FOXG1⁺ NPC model, we identified insulin/IGF signaling as the top enriched pathway. Notably, while IGF1R and IGF2R were highly expressed, IGF1 ligand was absent, suggesting that progenitors rely on external sources of IGF1. In rosette-derived neural aggregates, we discovered epithelial-like niche domains expressing IGF1 in direct contact with IGF1R⁺ FOXG1⁺ progenitors, defining a previously uncharacterized paracrine signaling architecture in early forebrain development.

Functionally, IGF1–IGF1R signaling activated PI3K–AKT–mTOR and MEK–ERK pathways to promote progenitor proliferation, clonal expansion, and vertical tissue growth. Genetic and pharmacological perturbations, along with BrdU incorporation and qPCR, showed that IGF1 modestly reinforced transcription of key identity genes (*FOXG1*, *PAX6*, *HES1*). In parallel, ribosome profiling, polysome analysis, and 5′UTR luciferase assays revealed that mTOR activation selectively enhanced translation of biosynthetic and neurodevelopmental transcripts, including *GSX1*—a ventral patterning gene implicated in interneuron specification and autism risk^22, 23^.

Together, these findings define a multilayered regulatory system in which niche-derived IGF1 coordinates transcriptional and translational programs via mTOR to sustain human forebrain progenitor fate and identity. This work establishes a new framework for understanding growth factor–niche interactions in cortical development and highlights selective translational control as a key mechanism linking extracellular cues to progenitor resilience and vulnerability^24^.

## Results

### Derivation of Human FOXG1⁺ Forebrain Neural Progenitors Reveals Niche-Dependent IGF1 Signaling Potential

To establish a human forebrain neural progenitor model, we derived FOXG1⁺ NPCs from human pluripotent stem cells using our previously published rosette-derived neural aggregate (RONA) culture system, which combines 3D embryoid body formation, 2D rosette induction, and 3D rosette-derived neural aggregation^25^ (Figure 1A–B). Immunofluorescence confirmed robust co-expression of FOXG1 and Nestin, canonical markers of forebrain identity and neural progenitors, respectively (Figure 1B).

**Figure 1.**
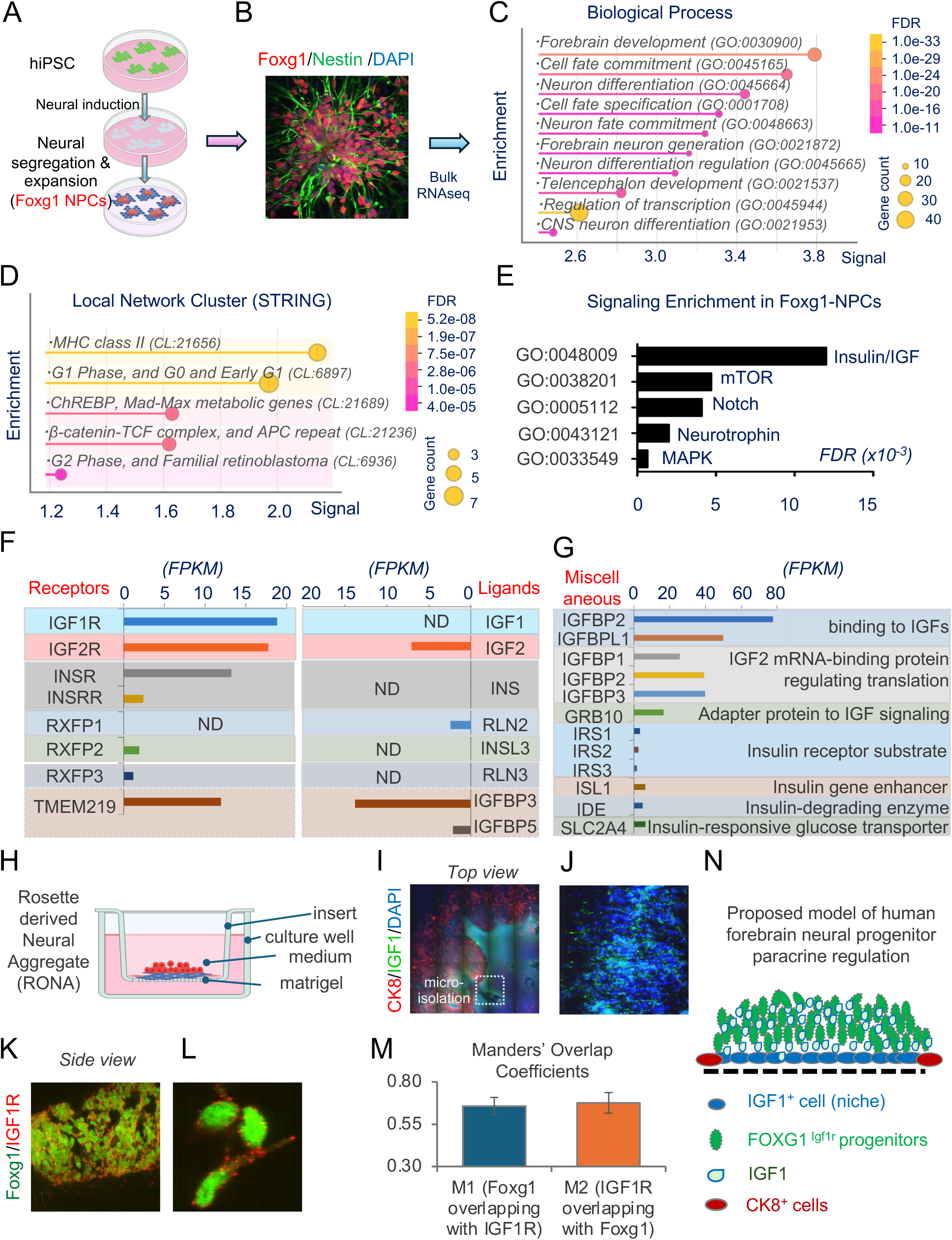
Characterization of human FOXG1⁺ neural progenitor cells and niche-dependent IGF1 signaling in rosette-derived neural aggregates. (**A-B**) Schematic illustrating derivation of FOXG1⁺ neural progenitor cells (NPCs) from human pluripotent stem cells using a 2D culture system. Immunostaining confirms co-expression of FOXG1 (red) and Nestin (green), with DAPI (blue) marking nuclei. (**C**) Gene Ontology (GO) enrichment analysis of FOXG1⁺ NPCs highlights processes including forebrain development, neuronal differentiation, and cell fate commitment. Dot size reflects the number of associated genes; color indicates false discovery rate (FDR). (**D**) STRING network analysis identifies enriched clusters involved in immune signaling, cell cycle regulation, and metabolic networks. (**E**) Pathway enrichment analysis shows strong activation of Insulin/IGF, mTOR, Notch, Neurotrophin, and MAPK signaling pathways. (**F**) Expression analysis reveals high levels of IGF1R and IGF2R, absent IGF1 ligand, and selective expression of Relaxin receptors, suggesting reliance on paracrine IGF1 signaling. (**G**) Expression of IGFBP2, IGFBPL1, IRS1–3, and regulators of IGF bioavailability and insulin signaling is detected, favoring IGF axis competence. (**H**) Schematic of rosette-derived neural aggregate (RONA) culture system showing 3D tissue organization supported by Matrigel and nutrient exchange. (**I-L**) Immunostaining of RONAs reveals central IGF1 expression (green), peripheral CK8⁺ epithelial cells (red), and FOXG1⁺ NPCs co-expressing IGF1R at the plasma membrane. (**M**) Quantitative co-localization analysis shows substantial overlap between FOXG1 and IGF1R (Manders’ coefficients ∼0.6). (**N**) Working model proposing that IGF1⁺ niche cells secrete IGF1 to regulate neighboring FOXG1⁺ progenitors in RONAs.

Transcriptomic profiling revealed that FOXG1⁺ NPCs were strongly enriched for forebrain development, cell fate commitment, neural differentiation, and telencephalon development pathways (Figure 1C and Table 1). STRING (Search Tool for the Retrieval of Interacting Genes/Proteins)^26^ network cluster analysis further highlighted enrichment in modules regulating cell cycle progression (G1/G2 phases), metabolic regulation (ChREBP–Max network, β-catenin– TCF/APC complex), and immune signaling (MHC class II) (Figure 1D). These modules indicated active regulation of proliferative programs, metabolic processes, and immune responses within the progenitor pool.

Unbiased signaling pathway enrichment^26, 27^ identified Insulin/IGF signaling as the top enriched axis, alongside mTOR, Notch, Neurotrophin, and MAPK pathways (Figure 1E). Intriguingly, targeted Insulin/IGF signaling component expression analysis revealed robust IGF1R and IGF2R expression, but absent IGF1 ligand production, while IGF2 was highly expressed (Figure 1F). Similarly, insulin receptors (INSR, INSRR) were expressed without detectable insulin (INS) ligand, and relaxin family pathways were minimally active. These patterns suggest that FOXG1⁺ NPCs are transcriptionally poised for IGF signaling but depend on external (paracrine) sources of IGF1. These expression patterns suggest that although FOXG1⁺ NPCs are transcriptionally poised for IGF axis activation, the cells lack intrinsic IGF1 production, implying a requirement for paracrine IGF1 signaling. Complementary profiling of regulatory components showed high expression of IGF modulators such as IGFBP2 and IGFBPL1 and moderate levels of IRS1–3 (insulin receptor substrates), with relatively lower expression of classical insulin signaling adaptors (GRB10, ISL1, IDE, SLC2A4) (Figure 1G). Together, these findings collectively support a model in which human forebrain progenitors are intrinsically primed for IGF-mediated paracrine signaling (Table 2).

To model niche-dependent signaling, we established RONA cultures using Matrigel-coated permeable inserts (Figure 1H). Multiplex immunostaining revealed localized IGF1 expression within the central domains beneath rosette-derived neural aggregates, surrounded by CK8⁺ epithelial-like boundary layers (Figure 1I-L). FOXG1⁺ progenitors displayed surface enrichment of IGF1R, positioning them to receive paracrine IGF1 cues. Co-localization analysis confirmed substantial overlap between FOXG1 and IGF1R signals (Manders’ coefficients ∼0.6) (Figure 1M), supporting active ligand-receptor interactions. Further assessment of IGF1 signaling competence, immunostaining confirmed widespread membrane localization of IGF1R in Nestin⁺ neural progenitors, exhibiting high co-localization (Manders’ coefficients ∼0.7) (Figure S1A–B).

We propose a working model in which IGF1⁺ niche cells secrete IGF1 ligands that act on adjacent FOXG1⁺ progenitors to sustain proliferation and neurogenic potential (Figure 1N). This niche-dependent mechanism establishes a novel paradigm of forebrain progenitor regulation distinct from canonical models based on intrinsic autocrine signaling.

Transcriptome-wide analysis of FOXG1⁺ NPCs identified expression of 515 transcription factors (TFs, FPKM>1), representing approximately 31% of the 1,639 high-confidence human TFs cataloged by Lambert et al^28^, these FOXG1⁺ NPC TFs were grouped into 15 clusters by k-means clustering^26^. The main Cluster 1 included 54 forebrain-enriched TFs (e.g., SOX2, FOXG1, SOX11, PAX6, OTX2) (Figure S1C–D, Table 3). Transcriptional network analysis revealed tightly interconnected hubs centered on FOXG1, PAX6, SOX2, and NEUROG2, suggesting coordinated regulation of forebrain progenitor identity (Figure S1D), with GO terms involved in regionalization, dorsal–ventral patterning, and G1 phase cell cycle regulation (Figure S1E-F). Selective expression of IGF modulators (CPE, EIF4EBP2, P4HB) and absence of IGF1, IGFBP1–5, and INSR (Figure S1G) reinforce the conclusion that FOXG1⁺ NPCs rely on external IGF1 inputs to activate IGF1R–mTOR signaling—revealing a niche-dependent regulatory framework essential for human forebrain development.

### Paracrine IGF1–IGF1R Signaling Promotes Clonal Expansion and Maintenance of Human Forebrain Neural Progenitors

Building upon the identification of niche-dependent IGF1 signaling, we next investigated the functional consequences of IGF1–IGF1R axis activation in human forebrain progenitors. Using the RONA culture system, we examined how manipulation of IGF1 availability affects progenitor expansion and tissue architecture (Figure 2).

**Figure 2.**
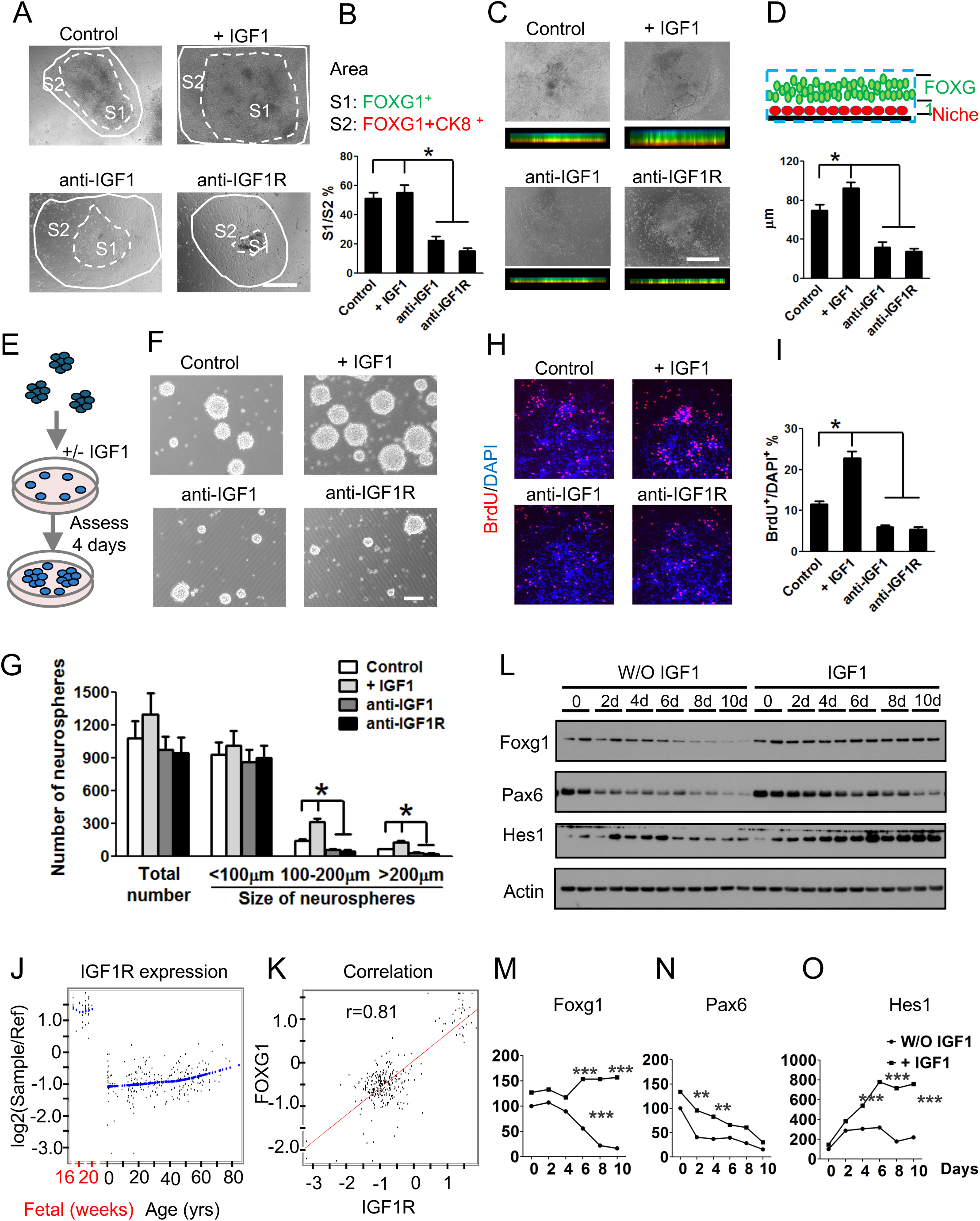
Paracrine IGF1–IGF1R signaling promotes clonal expansion and maintenance of human forebrain neural progenitors. **(A-B)** Brightfield imaging of RONAs shows regionalized progenitor (S1) and niche (S2) zones under control, IGF1 supplementation, IGF1 neutralization, and IGF1R blockade conditions. IGF1 maintains or enhances the S1/S2 ratio, while IGF1 or IGF1R blockade significantly reduces it (*p < 0.05). **(C-D)** Top and side-view images reveal that IGF1 promotes vertical tissue expansion, whereas IGF1 or IGF1R neutralization leads to thinner aggregates. **(E)** Quantification of aggregate thickness shows IGF1 significantly increases vertical growth compared to control, while neutralization treatments reduce it. **(F-G)** Neurosphere assays demonstrate that IGF1 enhances the size but not the total number of neurospheres, supporting its role in promoting clonal expansion. **(H-I)** BrdU incorporation assays show that IGF1 increases proliferation of FOXG1⁺ NPCs, while IGF1 or IGF1R blockade decreases BrdU⁺ cell fractions (*p < 0.05). **(J-K)** Developmental analysis reveals high IGF1R expression during early human fetal brain stages, correlating strongly with FOXG1 expression (r = 0.81). **(L-O)** Western blot and densitometry analyses demonstrate that IGF1 supplementation sustains FOXG1, Pax6, and Hes1 expression over time, supporting progenitor maintenance and survival.

Brightfield imaging revealed distinct structural zones within aggregates: a central progenitor-enriched S1 zone and a surrounding niche-enriched S2 zone. Under control conditions, the S1/S2 ratio remained stable, reflecting balanced progenitor maintenance (Figure 2A-B). Supplementation with recombinant IGF1 preserved or slightly enhanced the S1/S2 ratio, while neutralization of IGF1 or IGF1R markedly reduced it (*p < 0.05), indicating impaired progenitor expansion. Side-view imaging demonstrated that IGF1 treatment promoted vertical expansion and tissue thickness, whereas IGF1 or IGF1R blockade significantly reduced aggregate thickness (Figure 2C-D). These findings suggest that paracrine IGF1 not only sustains radial expansion but also supports vertical growth of progenitor structures.

Low-density neurosphere assays further revealed that while total sphere numbers remained comparable, IGF1 supplementation significantly increased the formation of larger neurospheres (>100 µm and >200 µm diameter), whereas IGF1 or IGF1R neutralization markedly reduced this capacity (*p < 0.05) (Figure 2E-G). Thus, IGF1 promotes clonal proliferative expansion rather than initial sphere formation. BrdU incorporation assays confirmed that IGF1 stimulation increased the fraction of proliferating progenitors, while IGF1 or IGF1R blockade significantly diminished proliferation rates (Figure 2H-I).

Analysis of human brain datasets^29^ revealed dynamic regulation of IGF1R expression during development, with peak IGF1R levels coinciding with maximal FOXG1 expression (Figure 2J-K). Scatter plot analysis showed a strong positive correlation (r = 0.81) between IGF1R and FOXG1, suggesting co-regulation of forebrain patterning and growth factor responsiveness.

Western blotting of neural progenitor markers across a 10-day culture timeline demonstrated that IGF1 supplementation preserved FOXG1 and Pax6 expression and enhanced HES1 levels, delaying differentiation-associated decline and supporting survival pathways (Figure 2L-O).

Immunostaining corroborated these findings: IGF1 increased Ki67⁺ cycling cells, while IGF1 or IGF1R blockade reduced the proliferative fraction (*p < 0.05), without affecting Caspase-3– positive apoptosis (Figure S2A–D).

To genetically validate these observations, we performed shRNA-mediated knockdown of IGF1R in FOXG1⁺ progenitors. Knockdown (especially with shRNA2) efficiently suppressed IGF1R protein levels (Figure S2E–F), resulting in reduced S1/S2 area ratios, impaired vertical growth (Figure S2G–J), and decreased BrdU incorporation (Figure S2K–L). Although neurosphere counts remained unchanged, sphere sizes were markedly diminished (Figure S2M–N). Immunostaining showed fewer Nestin⁺/Ki67⁺ progenitors after IGF1R depletion (Figure S2O–P). Finally, qPCR confirmed dynamic regulation of FOXG1, PAX6, and HES1 transcripts by IGF1: withdrawal reduced their expression, whereas IGF1 supplementation restored levels within 1 hour and further increased expression at 2 hours (Figure S2Q–T), consistent with sustained FOXG1 and Pax6 levels and upregulation of HES1 (Figure 2L–O).

Together, these findings establish that paracrine IGF1–IGF1R signaling is indispensable for sustaining human forebrain neural progenitor proliferation, clonal expansion, vertical tissue growth, and transcriptional maintenance.

### IGF1-IGF1R Signaling Activates ERK and Akt Pathways and Promotes Human Forebrain Neural Progenitor Proliferation Through PI3K–AKT–mTOR and MEK/ERK Cascades

To delineate downstream pathways engaged by IGF1–IGF1R signaling in human FOXG1⁺ neural progenitors, we examined the activation dynamics of ERK and Akt following IGF1 stimulation (Figure 3A-F). Western blot analysis revealed robust induction of ERK phosphorylation within 10 minutes, sustained through 60 minutes, without changes in total ERK levels (Figure 3A-B). Similarly, Akt phosphorylation increased rapidly upon IGF1 treatment, peaking at 10–60 minutes, while total Akt levels modestly declined over time, reflecting dynamic signaling regulation (Figure 3D-E).

**Figure 3.**
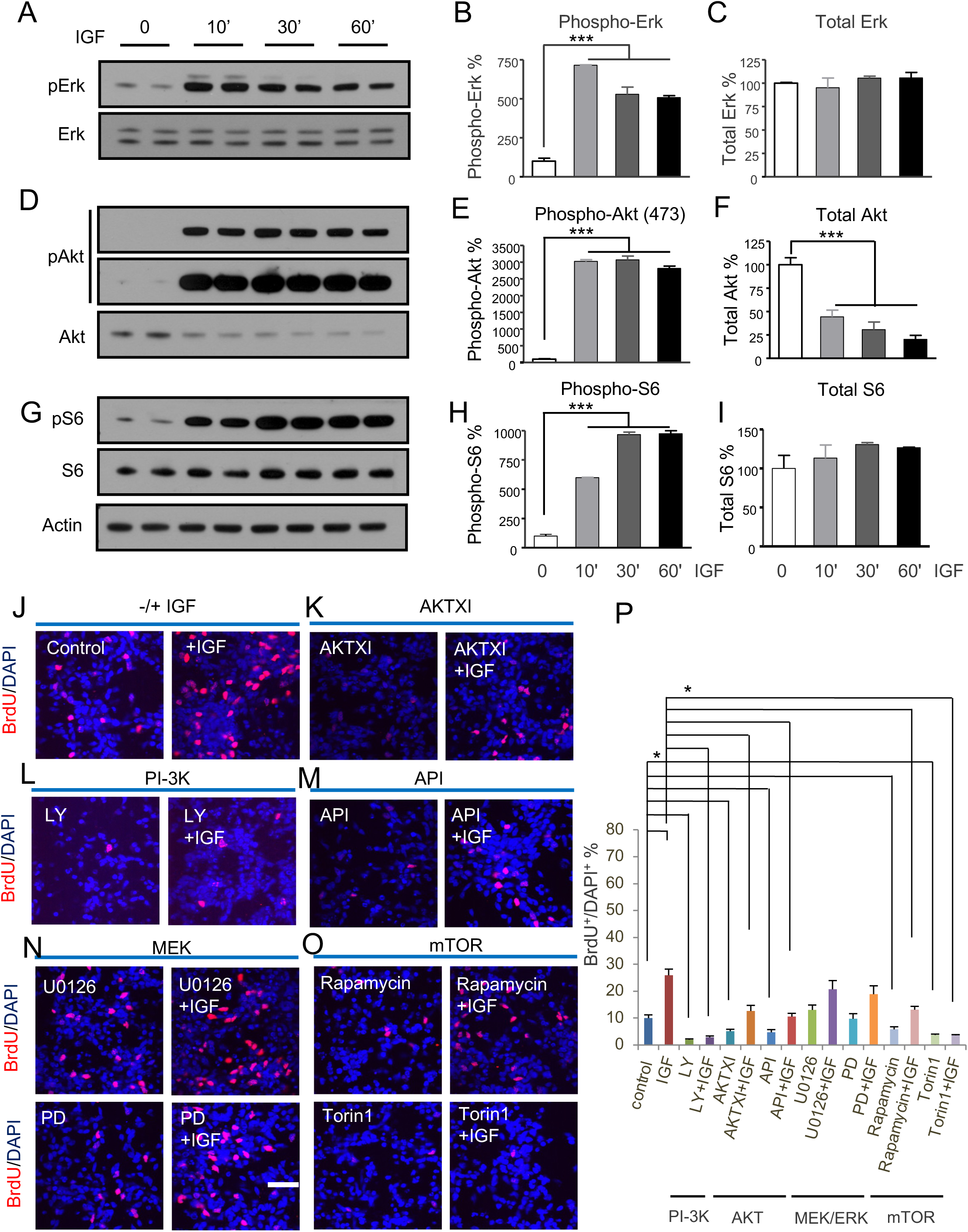
IGF1–IGF1R signaling activates ERK and Akt pathways and promotes forebrain neural progenitor proliferation through PI3K–AKT–mTOR and MEK/ERK cascades. **(A-C)** Western blot analysis shows that IGF1 rapidly and robustly induces ERK phosphorylation within 10 minutes, sustained up to 60 minutes. Total ERK levels remain stable. **(D-F)** IGF1 stimulation also induces Akt phosphorylation, peaking between 10–60 minutes, while total Akt levels modestly decline. **(G-I)** IGF1 enhances phosphorylation of S6, a downstream mTORC1 target, with peak activation at 30–60 minutes and stable total S6 levels. **(J-P)** BrdU incorporation assays combined with pathway-specific inhibitors demonstrate that PI3K, AKT, and mTOR pathways are essential for IGF1-driven proliferation. MEK/ERK inhibition partially reduces proliferation, indicating a secondary but supportive role.

Assessment of mTOR pathway activation showed that IGF1 stimulation markedly elevated phosphorylation of S6, a downstream mTORC target, reaching maximal activation by 30 minutes and remaining elevated at 60 minutes, with stable total S6 levels (Figure 3G-H).

To functionally dissect the role of these pathways in proliferation, BrdU incorporation assays were performed under IGF1 stimulation with selective pathway inhibitors (Figure 3J-P). IGF1 treatment significantly enhanced proliferation compared to controls. Inhibition of PI3K (LY294002, API), AKT (AKTXI), or mTOR (Rapamycin, Torin1) markedly suppressed IGF1-induced proliferation, underscoring the critical role of the PI3K–AKT–mTOR axis. MEK/ERK inhibition (U0126, PD032591) partially reduced proliferation, indicating a contributory but non-essential role.

These findings establish that IGF1–IGF1R signaling orchestrates forebrain progenitor proliferation primarily through PI3K–AKT–mTOR activation, with supportive input from MEK/ERK pathways.

### mTOR-4EBP1 Axis Is Required for IGF1-Induced Translational Activation and Neural Progenitor Proliferation

To further define mechanisms downstream of mTOR activation, we investigated the roles of mTOR and 4EBP1 in IGF1-mediated translational control (Figure S3). A paradigm of acute IGF1 stimulation following serum/growth factor deprivation was used to dissect signaling dynamics (Figure S3A-B). Western blotting confirmed that IGF1 enhanced phosphorylation of ERK, ribosomal protein S6 (RPS6), and 4EBP1, while mTOR inhibition with Torin1 selectively suppressed phosphorylation of S6 and 4EBP1 without affecting ERK activation (Figure S3C-F).

BrdU incorporation assays demonstrated that mTOR knockdown significantly suppressed proliferation despite IGF1 stimulation, indicating that mTOR activity is essential for IGF1-driven progenitor expansion (Figure S3G-H). In contrast, 4EBP1 knockdown paradoxically enhanced proliferation, suggesting that 4EBP1 serves as a negative regulator downstream of mTOR.

Together, these results define an IGF1–mTOR–4EBP1 signaling axis that governs global translational output, proliferation, and biosynthetic programs in human forebrain neural progenitors, providing new insights into how extrinsic IGF1 cues orchestrate intrinsic developmental pathways.

### IGF1 Orchestrates Regional Transcription, mTOR-Dependent Translation, Metabolic Pathways, and Genome integrity to Regulate Human Neural Progenitor Fate

To evaluate whether IGF1 signaling coordinates regional identity or maintains global transcriptional programs, we analyzed expression of key neurodevelopmental transcription factors following IGF1 modulation (Figure S4). IGF1 stimulation upregulated several ventral forebrain markers (e.g., DLX1, GSX2, ASCL1), although only DLX2 reached statistical significance (p = 0.0015). IGF1 withdrawal led to rapid downregulation of these genes. Similarly, progenitor markers such as SOX1 and OLIG2 showed an upward trend with IGF1 treatment, with significant upregulation observed for HES1 and SOX2 (p = 0.0473). In contrast, dorsal (EMX1/2) and posterior (HOXB4, KROX20) markers exhibited moderate or variable responses (Figure S4A–L). These findings suggest that while IGF1 modestly reinforces regional transcriptional programs, it does not drive broad transcriptional re-patterning. However, dynamic regulation of FOXG1, PAX6, and HES1 transcripts by IGF1 (Figure S2Q–T), along with sustained FOXG1 and PAX6 protein levels and upregulation of HES1 (Figure 2L–O), underscores IGF1’s role in maintaining core forebrain progenitor programs.

To dissect the downstream functional outputs of IGF1–mTOR signaling, we performed ribosome profiling (Ribo-seq)^24^ and polysome analysis to assess changes in translational regulation in FOXG1⁺ NPCs (Figure 4A). Unlike prior studies conducted under nutrient-rich conditions, our IGF1-deprivation and rescue approach enabled a focused analysis of acute, IGF1-induced, mTOR-dependent translational responses (Figure S4M). Ribosome footprinting confirmed strong ribosome accumulation at translation start sites following IGF1 stimulation, indicating efficient initiation of translation (Figure S4N). Cumulative distribution analysis revealed that IGF1 stimulation significantly enhanced global translation efficiency compared to baseline, mTOR inhibition with Torin1 moderately attenuated these effects (Figure 4B-C, Table 4). Analysis of known mTOR target transcripts demonstrated that IGF1-induced translation of these genes was highly sensitive to Torin1, confirming that mTOR activation is required for efficient translational upregulation. Pathway enrichment analysis of transcripts with increased translation efficiency identified ribosome biogenesis as the top pathway, followed by oxidative phosphorylation, proteasome function, and neurodegenerative disease-associated processes such as Parkinson’s and Huntington’s disease (Figure 4D). Gene Ontology enrichment analyses further revealed that IGF1 stimulation upregulated translation-related biosynthetic programs, including cytoplasmic translation, ribonucleoprotein complex assembly, translational elongation, and ribosome assembly (Figure 4E). In addition to biosynthesis, IGF1-enhanced translation programs prominently involved energy metabolism pathways such as oxidative phosphorylation and mitochondrial organization (Figure 4F-H), as well as proteostasis mechanisms including ubiquitin-mediated protein catabolism and multivesicular body trafficking (Figure 4G).

**Figure 4.**
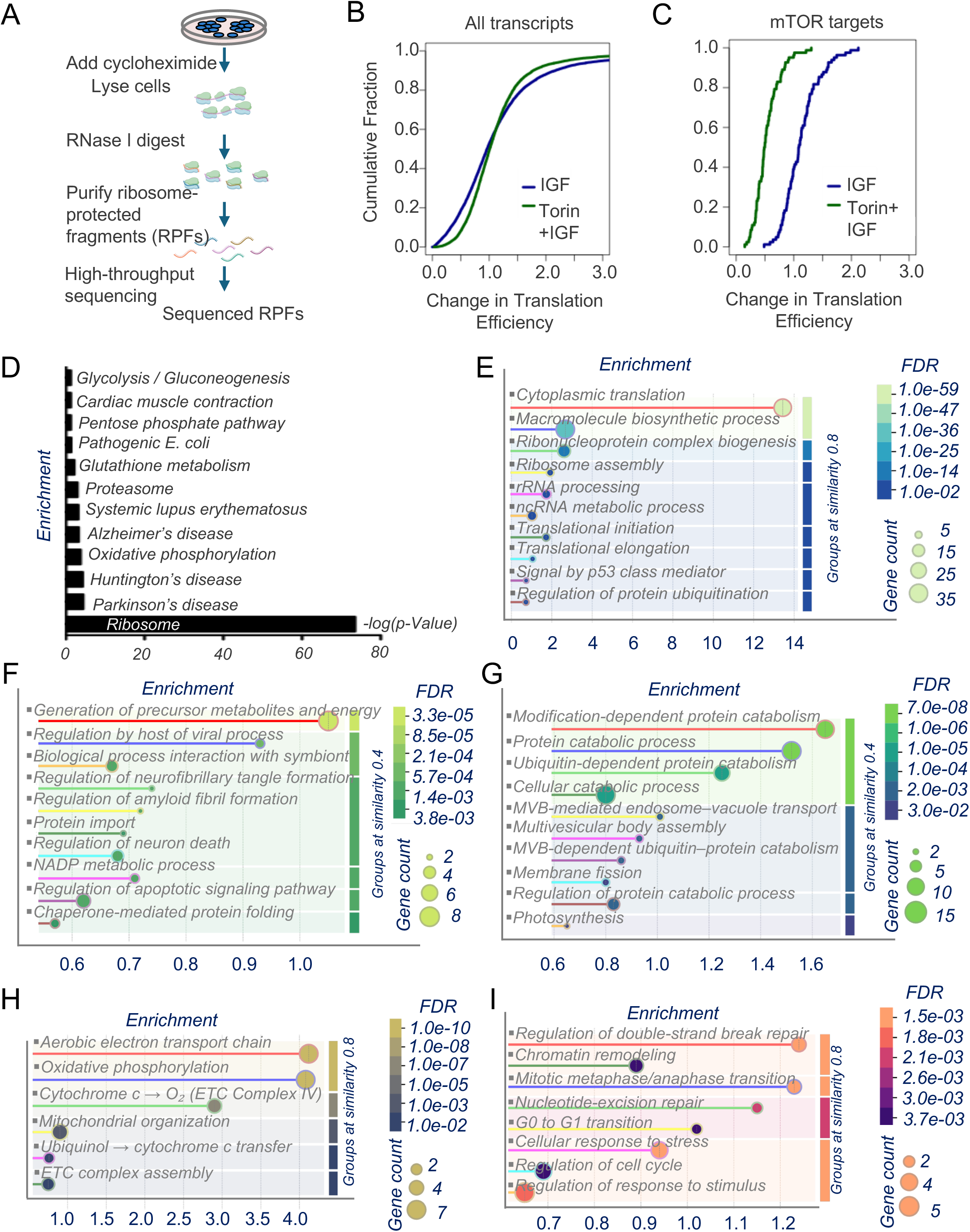
IGF1–mTOR Axis regulates neural progenitor translational programs governing biosynthesis, energy metabolism, proteostasis, and genome stability. **(A)** Workflow illustrating ribosome profiling (Ribo-Seq) in cultured FOXG1⁺ neural progenitors, including cycloheximide treatment, ribosome-protected fragment (RPF) isolation, and sequencing. **(B-C)** Cumulative distribution plots show that IGF1 broadly enhances translation efficiency transcriptome-wide, while Torin1 attenuates this effect, particularly for known mTOR targets. **(D)** KEGG pathway enrichment identifies ribosome biogenesis, oxidative phosphorylation, proteasome function, and neurodegenerative pathways (e.g., Parkinson’s, Huntington’s) among IGF1-upregulated translational targets. **(E-F)** GO biological processes enriched among IGF1-upregulated transcripts include cytoplasmic translation, ribonucleoprotein biogenesis, and macromolecule biosynthesis. **(G)** IGF1-responsive translational programs prominently involve proteostasis mechanisms such as ubiquitin-dependent protein catabolism and multivesicular body transport. **(H)** IGF1 stimulation enhances translation of mitochondrial pathways, including the electron transport chain and oxidative phosphorylation. **(I)** IGF1 promotes translation of genes involved in genome stability, including DNA repair, chromatin remodeling, and cell cycle checkpoints.

Additionally, IGF1-responsive translational programs were enriched for processes related to genome integrity, including DNA repair, chromatin remodeling, mitotic regulation, and cellular stress responses (Figure 4I). Supporting these findings, GO enrichment also highlighted mRNA stabilization, splicing, and nucleobase metabolism (Figure S4O). At the structural level, regulated transcripts were linked to paranodal junctions and L1-ankyrin interactions, suggesting effects on cytoskeletal organization (Figure S4P). Finally, metabolic pathway analysis revealed enrichment in cysteine and methionine metabolism and broader metabolic networks (Figure S4Q), reinforcing the role of IGF1–mTOR signaling in coordinating biosynthetic, energetic, and proteostatic demands essential for progenitor maintenance.

These findings demonstrate that IGF1–mTOR signaling orchestrates a multilayered enhancement of biosynthesis, energy metabolism, proteostasis, and genome integrity— processes essential for maintaining human forebrain progenitor homeostasis. Collectively, the results reveal that IGF1 integrates niche-derived cues with both regional transcriptional programs and global translational control, linking extrinsic growth factor availability to intrinsic developmental and metabolic regulation required for progenitor expansion and patterning.

### Selective Translational Control of Non-Canonical mTOR Targets Links IGF1 Signaling to Neurodevelopment and Progenitor Niche Regulation

While many mTOR-responsive mRNAs encode ribosomal proteins, elongation factors, and initiation factors as shown previously by Thoreen etc^30^. and Hsieh etc.^31^, we also identified transcripts with non-canonical or absent TOP motifs that exhibited increased translation upon IGF1 stimulation (Figure S5A–E). Pathway enrichment of these non-conventional targets revealed upregulation of neurodevelopmental processes—including nervous system development and axon guidance—and biosynthetic programs such as cap-dependent and cytoplasmic translation (Figure 5A).

**Figure 5.**
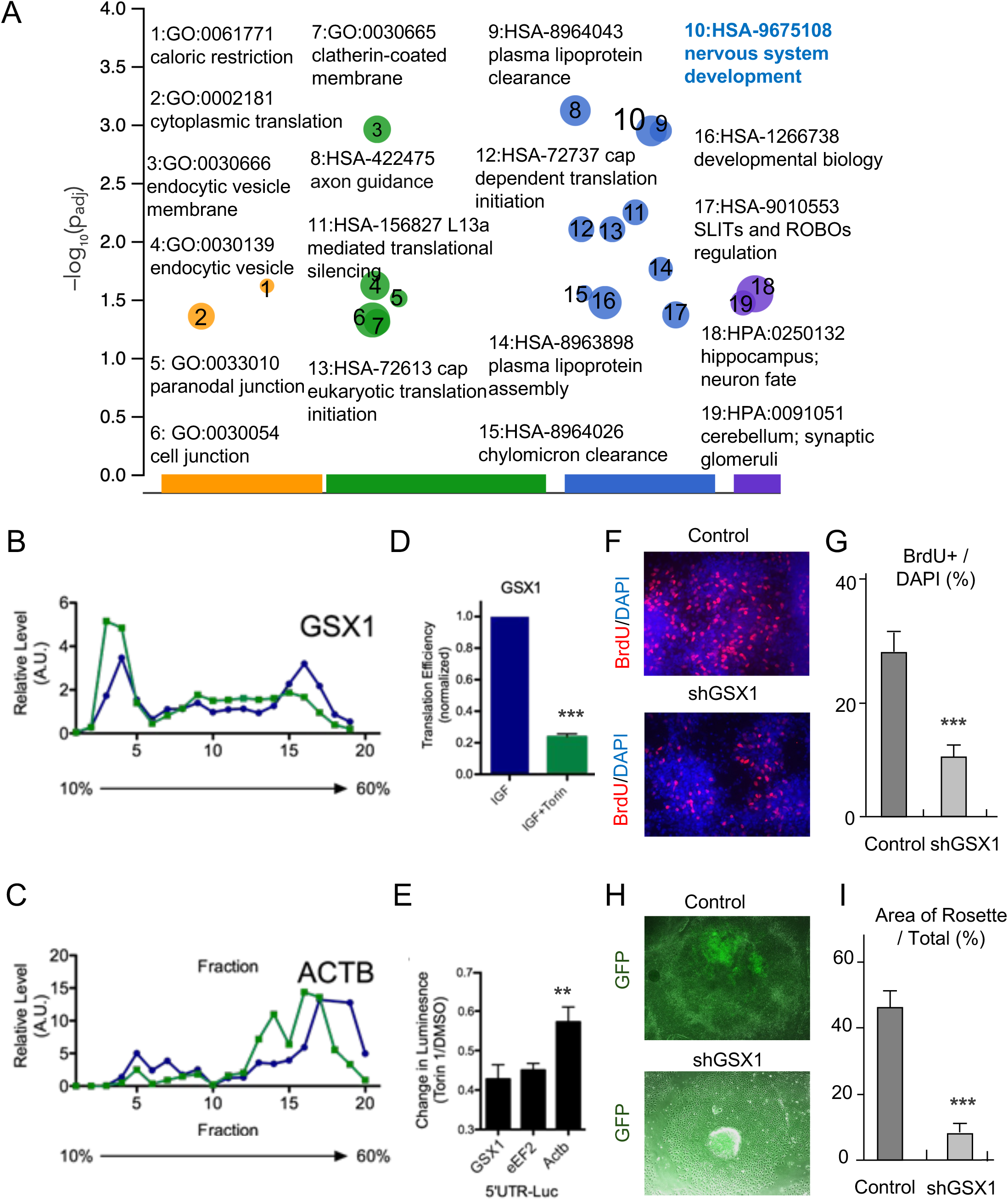
IGF1-driven translational programs in human neural progenitors are mediated by mTOR activation and regulated by 5’UTR-dependent mechanisms. **(A)** Gene Ontology and pathway enrichment analysis identifies translational programs regulated by IGF1, including nervous system development, axon guidance, ribosome biogenesis, cytoplasmic translation, lipoprotein metabolism, and neuronal maturation pathways (e.g., SLIT– ROBO signaling). **(B-C)** Polysome fractionation analysis shows that IGF1 promotes GSX1 and ACTB mRNA association with heavier polysome fractions, indicating enhanced translational engagement, while Torin1 treatment shifts transcripts toward lighter fractions. **(D)** Bar graph quantifying polysome-to-monosome ratios reveals that IGF1 significantly enhances global translational output, while mTOR inhibition (Torin1) suppresses this effect. **(E)** 5’UTR reporter assays demonstrate that luciferase reporters bearing the GSX1, eEF2, or ACTB 5’UTRs exhibit reduced translation upon Torin1 treatment, with GSX1 5’UTR showing the greatest sensitivity, indicating 5’UTR-dependent mTOR control. **(F-G)** Immunofluorescence images and quantification show that expression of a GSX1 5’UTR– luciferase reporter reduces BrdU incorporation compared to control reporters, suggesting translational repression impairs progenitor proliferation. **(H-I)** Brightfield and GFP imaging reveal that GSX1 5’UTR–GFP constructs significantly reduce GFP intensity compared to control, confirming strong 5’UTR-mediated translational repression affecting protein output in neural progenitor colonies.

To probe transcript-specific translational control, we combined polysome profiling with 5′UTR luciferase assays. IGF1 stimulation enhanced heavy polysome formation, while mTOR inhibition with Torin1 reduced polysome abundance, confirming mTOR’s role in global translational engagement (Figure S5A). GSX1, a neurodevelopmental transcription factor, emerged as a representative IGF1–mTOR target. IGF1 promoted GSX1 and ACTB mRNA association with heavy polysome fractions, indicating active translation, whereas Torin1 shifted them to lighter fractions (Figure 5B–C). IGF1 also increased the global polysome-to-monosome ratio, further supporting a boost in translational output (Figure 5D).

Luciferase reporter assays using the 5′UTRs of GSX1, eEF2, and ACTB confirmed 5′UTR-dependent translational regulation. Among them, the GSX1 5′UTR was most sensitive to mTOR inhibition, suggesting strong mTOR-dependent control (Figure 5E). Functionally, GSX1 knockdown in FOXG1⁺ progenitors reduced BrdU incorporation and impaired radial expansion of rosette structures, confirming its role in progenitor proliferation (Figure 5F–I).

To further elucidate how IGF1–mTOR–regulated transcripts lacking canonical or containing non-canonical TOP motifs shape cellular functions in FOXG1⁺ progenitors, we applied NetworkAnalyst and constructed protein–protein interaction (PPI) networks^32, 33^ based on IGF1-responsive translational targets (Figure S5B-G). PPI network analysis revealed that IGF1-regulated seed proteins, including ATL3, UBXN10, ATP6V0A1, COA3, EP400, GEN1, and ONECUT1, formed highly connected hubs involved in mitochondrial function, vesicle trafficking, transcriptional regulation, and neurodevelopment (Figure S5B). Focused networks highlighted that CPAMD8 and MMP17 were centrally involved in extracellular matrix remodeling and proteolytic regulation (Figure S5C), suggesting that IGF1 may influence progenitor niche architecture through matrix dynamics. NSMF-centered networks implicated IGF1-responsive translation in axon guidance and neuroendocrine signaling pathways, reflecting broad developmental roles (Figure S5D). SPING1 and C1orf122 networks suggested regulation of proteostasis, membrane trafficking, and chromatin remodeling (Figure S5E-F), linking IGF1 signaling to stress response and genome stability mechanisms. GSX1-centered networks implicated neural fate commitment (Figure S5G).

These results demonstrate that IGF1–mTOR signaling enhances translational output in human forebrain progenitors by selectively regulating key neurodevelopmental transcripts. This provides a mechanistic link between external niche-derived cues and the maintenance of progenitor proliferation and identity. Beyond individual gene targets, IGF1–mTOR–mediated translational reprogramming broadly remodels protein interaction networks, coordinating biosynthesis, energy metabolism, proteostasis, cytoskeletal dynamics, and developmental programs essential for progenitor maturation and neurodevelopmental resilience.

## Discussion

This study defines a multilayered regulatory network in human forebrain neural progenitor cells (NPCs), orchestrated by niche-derived IGF1 and mediated through mTOR signaling. By integrating transcriptomic and ribosome profiling with functional assays, we demonstrate that IGF1 sustains forebrain NPC proliferation, identity, and differentiation via coordinated transcriptional and translational control.

Transcriptomic analysis revealed insulin/IGF signaling as the most enriched pathway in FOXG1⁺ NPCs, with robust expression of IGF1R and IGF2R but absence of IGF1 ligand—indicating dependence on external IGF1. Our 3D RONA culture system revealed epithelial-like HNF3β⁺ (FOXA2⁺) niche domains expressing IGF1 adjacent to IGF1R⁺ progenitors, establishing a spatially organized, paracrine signaling architecture unique to early forebrain development.

Functionally, IGF1 promoted progenitor proliferation, vertical expansion, and neuroepithelial marker preservation, while inhibition of IGF1 or IGF1R impaired growth and transcriptional maintenance. At the molecular level, IGF1 reinforced key transcription factors (e.g., *FOXG1*, *PAX6*, *HES1*) and activated PI3K–AKT–mTOR and MEK–ERK pathways. Importantly, transcriptional regulators were not themselves translational targets, indicating a division of regulatory labor between transcriptional support and translational output.

At the molecular level, our results reveal a dual regulatory mechanism: IGF1 reinforces core transcriptional programs (e.g., FOXG1, PAX6, HES1) while also driving distinct, mTOR-dependent translational programs. Although IGF1 modestly modulates regional identity markers, it does not induce broad transcriptional re-patterning. Instead, IGF1 sustains key forebrain progenitor TFs that regulate neuroepithelial stability. Importantly, these transcriptional regulators are not themselves translational targets of IGF1–mTOR signaling, highlighting the mechanistic distinction between transcriptional reinforcement and translational reprogramming.

Ribosome profiling identified IGF1–mTOR–responsive transcripts enriched for neurodevelopmental, biosynthetic, and metabolic functions—many lacking canonical TOP motifs. These mTOR-dependent translational programs expand the biosynthetic and proteostatic capacity of NPCs, enabling sustained growth and differentiation.

A notable discovery was the identification of *GSX1* as a direct, mTOR-sensitive translational target. GSX1, a homeobox TF essential for ventral telencephalic patterning and interneuron lineage specification^23^, showed 5′UTR-dependent translational upregulation by IGF1.

Knockdown impaired progenitor proliferation and radial expansion, underscoring its role in progenitor competence. This finding adds a previously unrecognized layer of translational control to interneuron lineage specification and links niche-derived growth signals to autism-relevant circuit development. Given the co-occurrence of *GSX1* dysregulation, mTOR hyperactivity, and interneuron imbalance in autism spectrum disorders (ASDs), our work positions *GSX1* as a critical node linking external cues to neurodevelopmental vulnerability.

Beyond cell-intrinsic programs, IGF1-regulated transcripts formed protein interaction hubs involved in mitochondrial function, vesicular trafficking, proteostasis, and matrix remodeling— highlighting how IGF1–mTOR signaling also shapes progenitor–niche interactions. These insights reveal potentially distinct regulatory dynamics in human NPCs compared to rodent models, underscoring the value of human systems for interrogating species-specific neurodevelopmental mechanisms.

A key strength of our study is the use of developmentally authentic human FOXG1⁺ NPCs, which express a rich complement of transcription factors (∼31% of all high-confidence human TFs) organized into forebrain-relevant regulatory networks. Unlike prior studies using transformed cell lines or non-neural systems^30,31^ we employed an IGF1-deprivation and rescue strategy under serum-free conditions, enabling resolution of acute, ligand-specific mTOR-driven translational responses in a fate-restricted neural progenitor population.

In summary, our findings establish a paradigm in which paracrine IGF1–mTOR signaling integrates transcriptional and translational programs to maintain human forebrain NPC fate. This work provides a mechanistic framework for how growth factor niches guide cortical development and identifies selective translation as a critical regulatory axis. Future studies should expand on this by mapping IGF1–mTOR dynamics across developmental trajectories and examining how genetic perturbations of mTOR-sensitive transcripts influence progenitor function in ASDs and related disorders. Therapeutic modulation of this pathway in human brain organoids may ultimately inform strategies for neurodevelopmental intervention and brain repair.

### Study Limitations and Implications

While our study defines a niche-dependent IGF1–mTOR signaling axis that integrates transcriptional and translational regulation in human forebrain neural progenitors, several limitations should be acknowledged. First, although the RONA culture system generates patterned FOXG1⁺ NPCs and recapitulates localized paracrine interactions, it lacks vascularization and perfusion, limiting insights into systemic or long-range signaling cues that may modulate IGF1 responsiveness.

Second, our ribosome profiling focused on acute IGF1 rescue following deprivation. Long-term or graded IGF1 stimulation, or chronic mTOR modulation, may reveal additional layers of regulation—such as feedback inhibition, mTORC2 contributions, or dynamic changes in translational selectivity.

Third, while GSX1 knockdown impaired progenitor proliferation and structural expansion, its lineage-specific roles remain to be elucidated. Moreover, the relevance of IGF1–mTOR–GSX1 signaling to autism spectrum disorder (ASD) and other neurodevelopmental disorders remains correlative and warrants in vivo validation using humanized or chimeric brain models.

Despite these limitations, our findings demonstrate that early human FOXG1⁺ NPCs rely on paracrine IGF1 input to activate IGF1R–mTOR signaling—establishing a regulatory architecture distinct from classical autocrine models. This underscores the essential role of niche-derived IGF1 in sustaining progenitor identity and expansion. Importantly, the identification of non-canonical, mTOR-sensitive translational targets such as *GSX1* extends the known repertoire of mTOR-regulated processes beyond classical TOP-containing transcripts.

Our work also has implications for understanding TSC1-associated cortical pathology. TSC1 negatively regulates mTORC1, and its loss leads to cortical tuber formation—malformations that arise specifically during prenatal neurogenesis from dysregulated mTOR signaling in fate-restricted forebrain progenitors. Postnatal or adult mTOR dysregulation does not recapitulate this phenotype, emphasizing the critical timing of mTOR activity in cortical development. In this context, our FOXG1⁺ NPC model provides a human-relevant platform to dissect how growth factor–mTOR interactions intersect with TSC1 loss-of-function during this vulnerable developmental window.

Finally, by linking IGF1–mTOR signaling to the control of biosynthesis, metabolism, proteostasis, and genome integrity, our findings highlight this pathway’s central role in progenitor resilience. These insights inform the refinement of forebrain organoid systems and may guide translational strategies for treating mTOR-related neurodevelopmental disorders and enabling brain repair.

---

**Figure S1.**
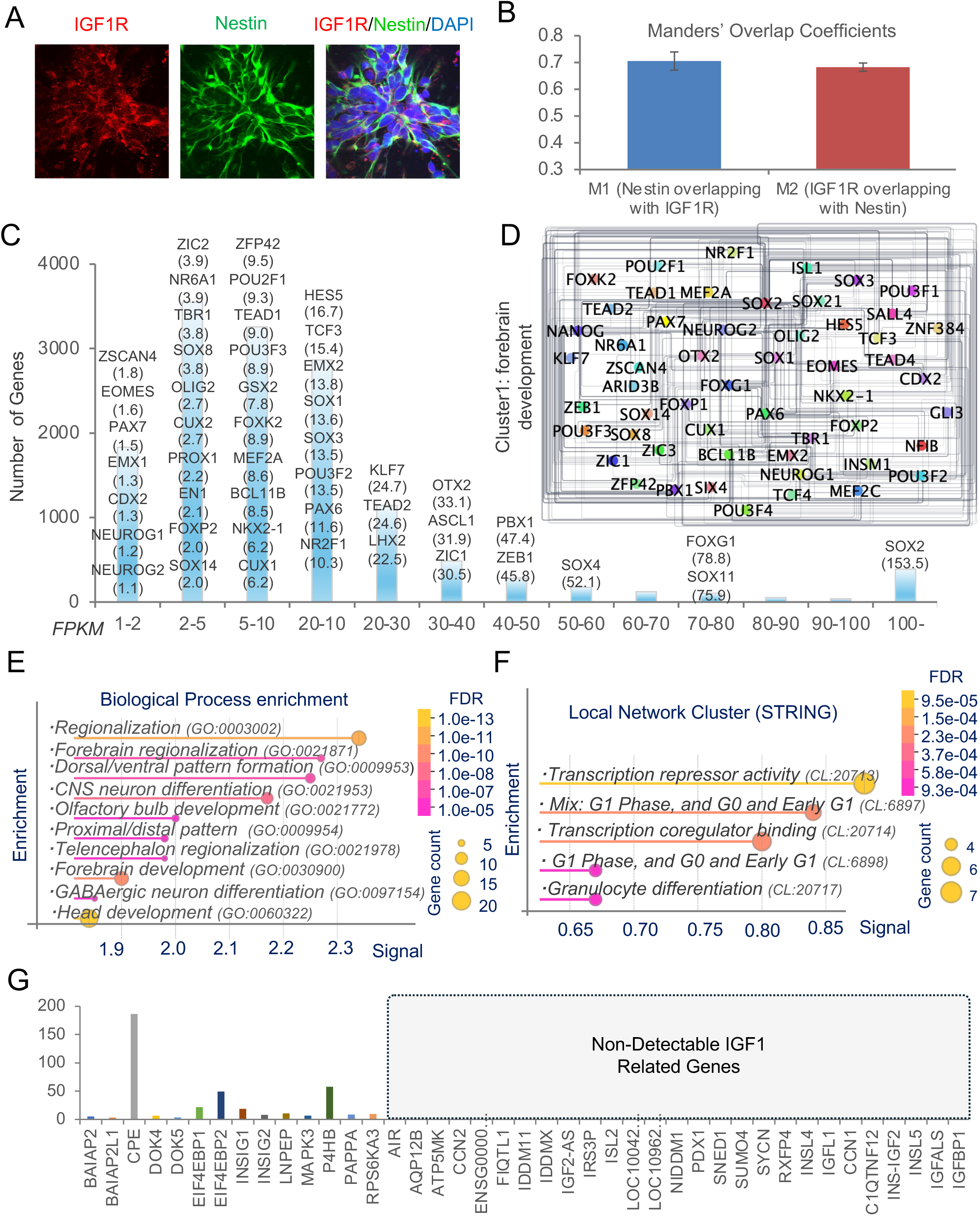
IGF1R localization and transcriptional programs sustaining forebrain identity in FOXG1⁺ neural progenitor cells. (**A-B**) Immunofluorescence shows IGF1R localization along Nestin⁺ progenitor membranes. High co-localization (Manders’ coefficients ∼0.7) confirms accessibility to IGF1. (**C**) Gene expression distribution in FOXG1⁺ NPCs. Highly expressed forebrain-associated transcription factors include SOX2, FOXG1, SOX11, and PAX6. (**D**) Forebrain transcriptional regulatory network mapping shows hubs around FOXG1, PAX6, SOX2, EMX2, and NEUROG2. (**E**) GO enrichment highlights processes including regionalization, dorsal-ventral patterning, and CNS neuron differentiation. (**F**) STRING cluster analysis reveals modules regulating transcriptional repression and cell cycle control. (**G**) Expression analysis shows selective detection of IGF1-modulatory genes (e.g., CPE, EIF4EBP2), but absent expression of classical IGF1 ligands or receptors, reinforcing the requirement for paracrine IGF1 input.

**Figure S2.**
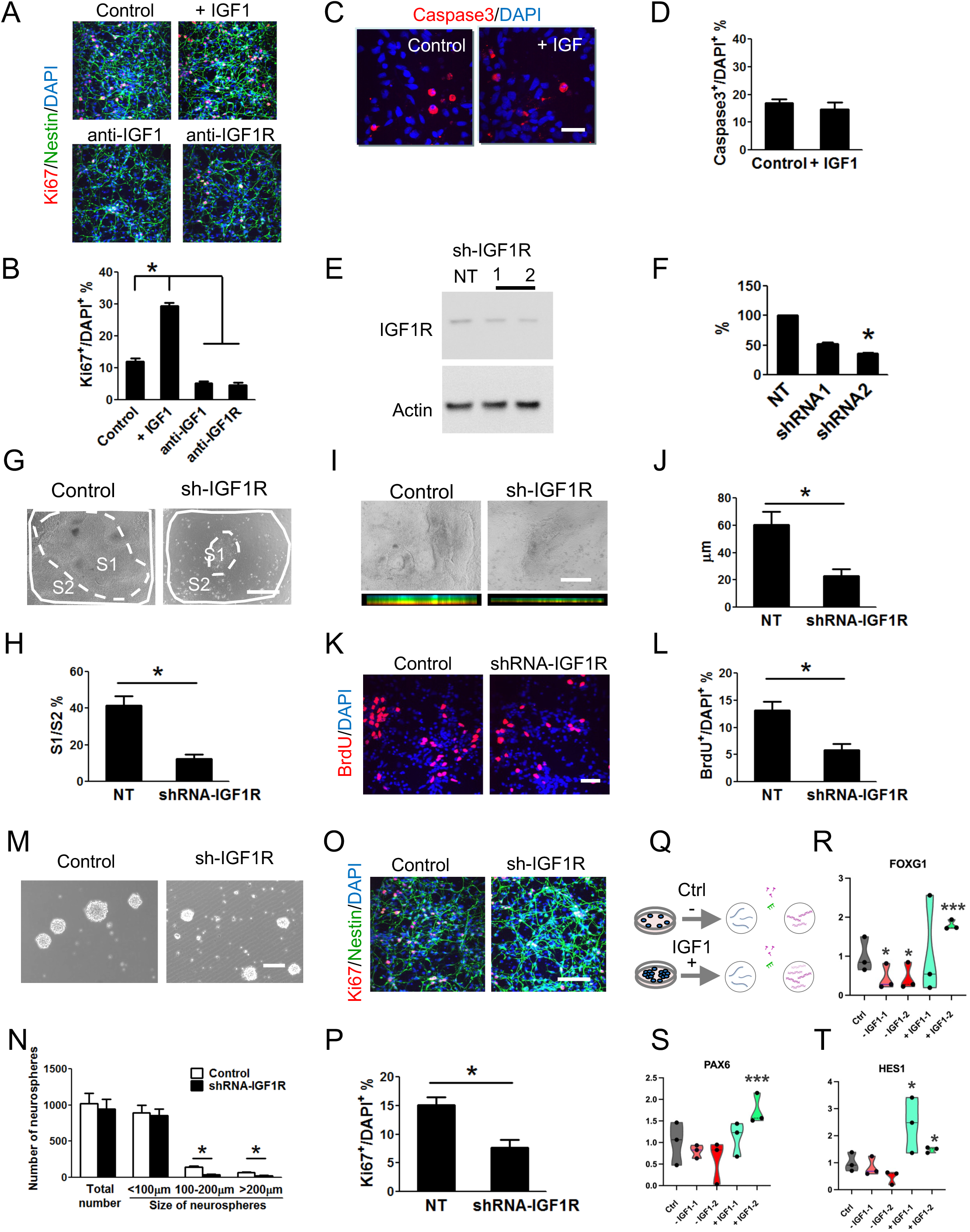
IGF1–IGF1R axis controls proliferation, expansion, and transcriptional identity of human forebrain neural progenitors. **(A-B)** Ki67 immunostaining shows that IGF1 enhances progenitor proliferation, whereas IGF1 or IGF1R blockade reduces Ki67⁺ fractions (*p < 0.05). **(C-D)** Caspase-3 staining indicates that IGF1 does not significantly alter apoptosis under basal conditions. **(E-F)** Western blot validation confirms efficient IGF1R knockdown using two independent shRNAs, with shRNA2 achieving stronger suppression. **(G-H)** Brightfield imaging shows that IGF1R knockdown reduces S1/S2 ratios, indicating impaired progenitor expansion. **(I-J)** Side-view imaging demonstrates that IGF1R knockdown results in thinner RONAs, confirming loss of vertical growth capacity. **(K-L)** BrdU incorporation assays show reduced S-phase proliferation upon IGF1R knockdown (*p < 0.05). **(M-N)** Neurosphere assays reveal that IGF1R knockdown decreases neurosphere size without affecting total neurosphere number. **(O-P)** Ki67 and Nestin co-staining confirms that IGF1R knockdown reduces the proportion of proliferating progenitors. **(Q-T)** Dynamic qPCR analysis shows that FOXG1, PAX6, and HES1 expression are rapidly downregulated upon IGF1 deprivation and upregulated with IGF1 supplementation, supporting IGF1-dependent maintenance of progenitor transcriptional identity.

**Figure S3.**
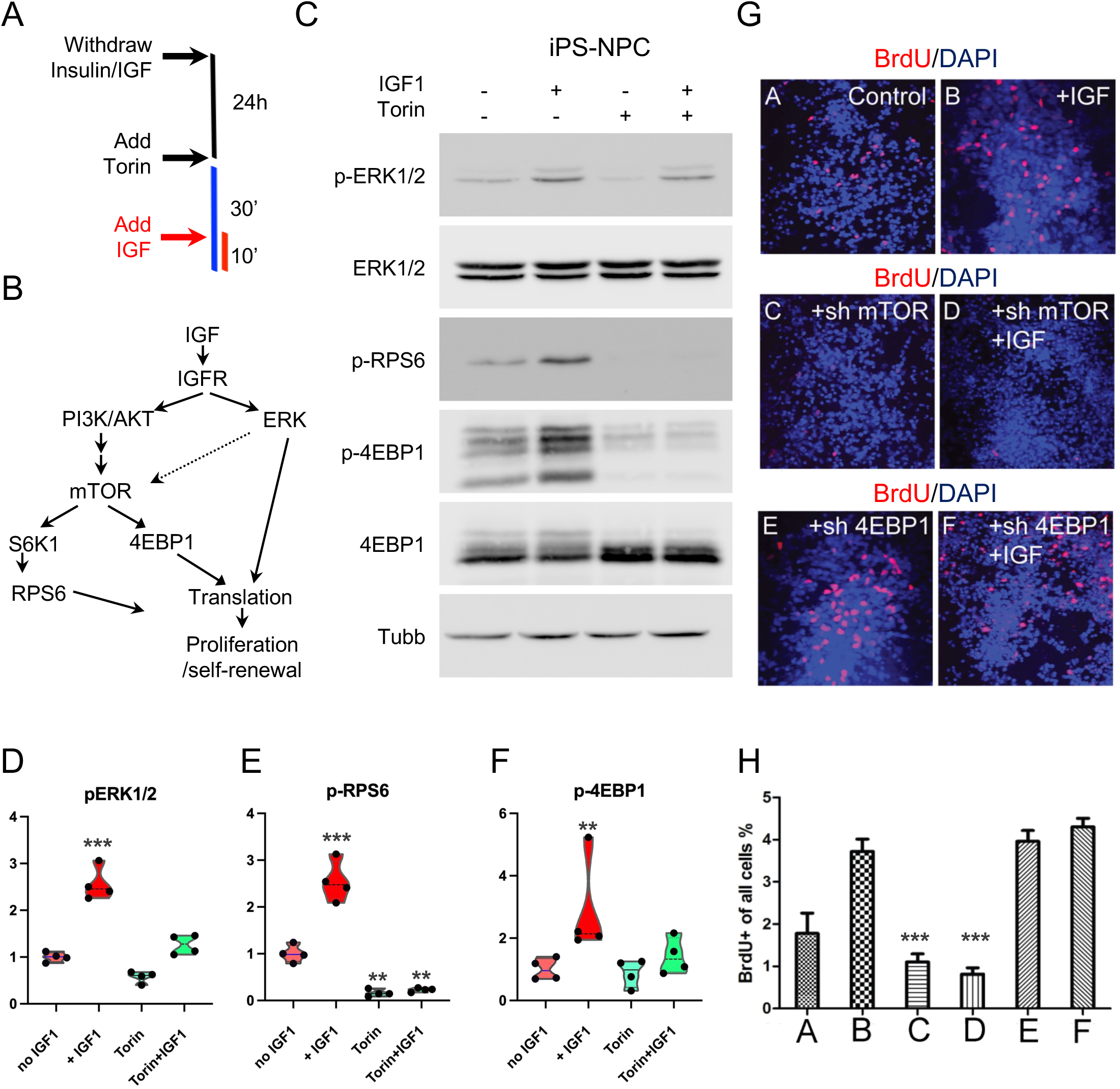
IGF1 Activates ERK and mTOR Signaling to Promote Proliferation in Human Forebrain Neural Progenitors via a 4EBP1-Dependent Mechanism. **(A-B)** Schematic showing experimental setup for acute IGF1 stimulation after growth factor deprivation. **(C-F)** Western blot analyses show that IGF1 enhances phosphorylation of ERK, RPS6, and 4EBP1. mTOR inhibition with Torin1 abolishes phosphorylation of RPS6 and 4EBP1 but does not affect ERK activation. **(G-H)** BrdU assays reveal that mTOR knockdown suppresses IGF1-driven proliferation, while 4EBP1 knockdown enhances proliferation, indicating that mTOR promotes growth while 4EBP1 constrains it.

**Figure S4.**
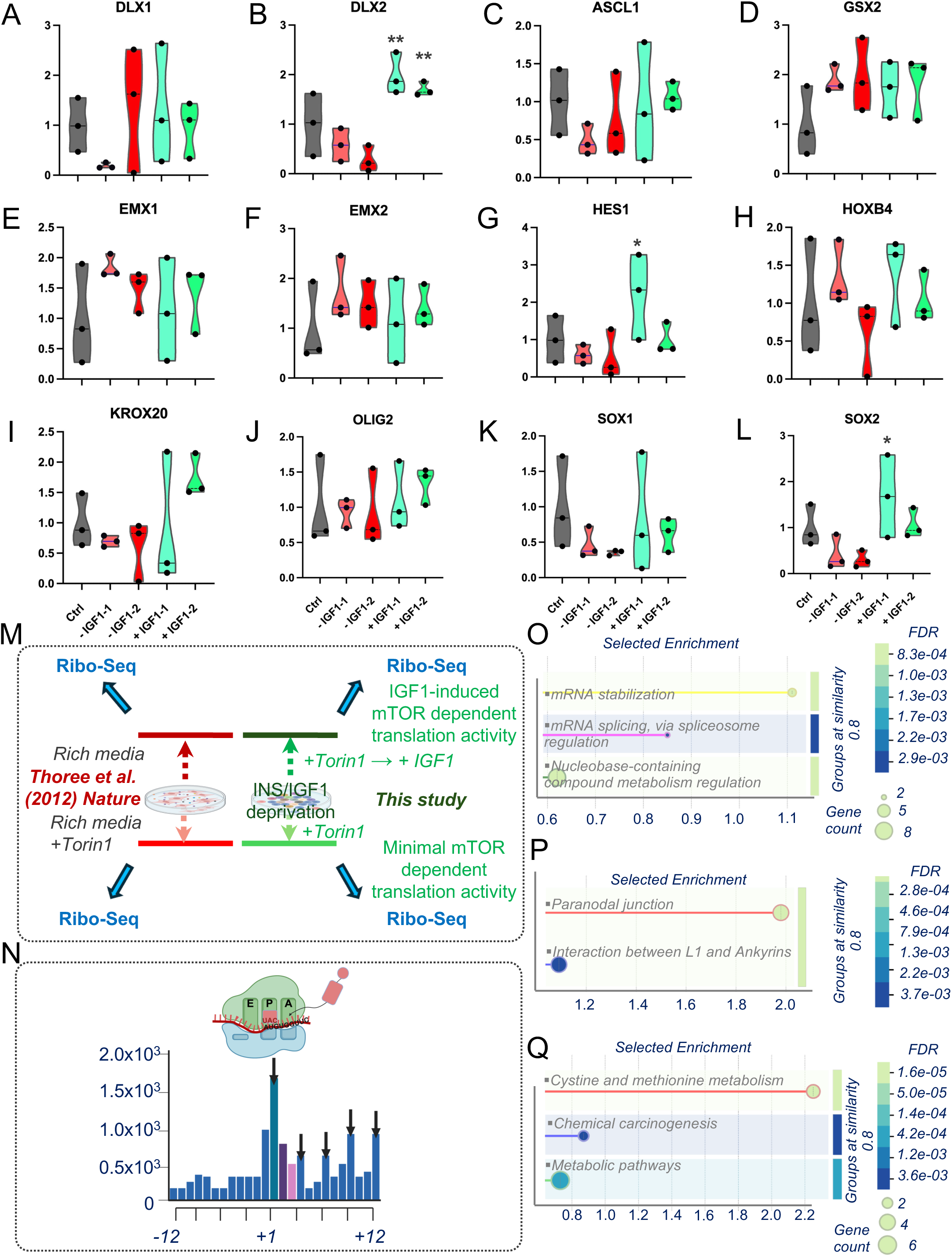
IGF1 orchestrates regional transcriptional programs, mTOR-dependent translation, and neurodevelopmental pathways in forebrain neural progenitors. **(A-L)** Violin plots show that IGF1 stimulation upregulates ventral forebrain markers (DLX1, DLX2, GSX2, ASCL1), Notch effector HES1, and neural progenitor markers (SOX2, SOX1, OLIG2), while dorsal markers (EMX1, EMX2) and posterior genes (HOXB4, KROX20) are less responsive. **(M)** Comparison of experimental designs: unlike prior nutrient-rich paradigms, our model uses IGF1 re-stimulation after deprivation, allowing acute dissection of IGF1-mTOR translational control. **(N)** Ribosome profiling reveals strong ribosome occupancy at translation start sites, confirming high precision of translation initiation capture. **(O)** GO enrichment highlights regulation of mRNA stabilization, splicing, and nucleobase metabolism among IGF1-responsive translationally upregulated transcripts. **(P)** Enriched cellular components include paranodal junctions and L1–ankyrin interactions, linking IGF1 translation targets to neurostructural development. **(Q)** Metabolic pathway enrichment shows IGF1 regulates cysteine/methionine metabolism, chemical carcinogenesis pathways, and broad metabolic processes at the translational level.

**Figure S5.**
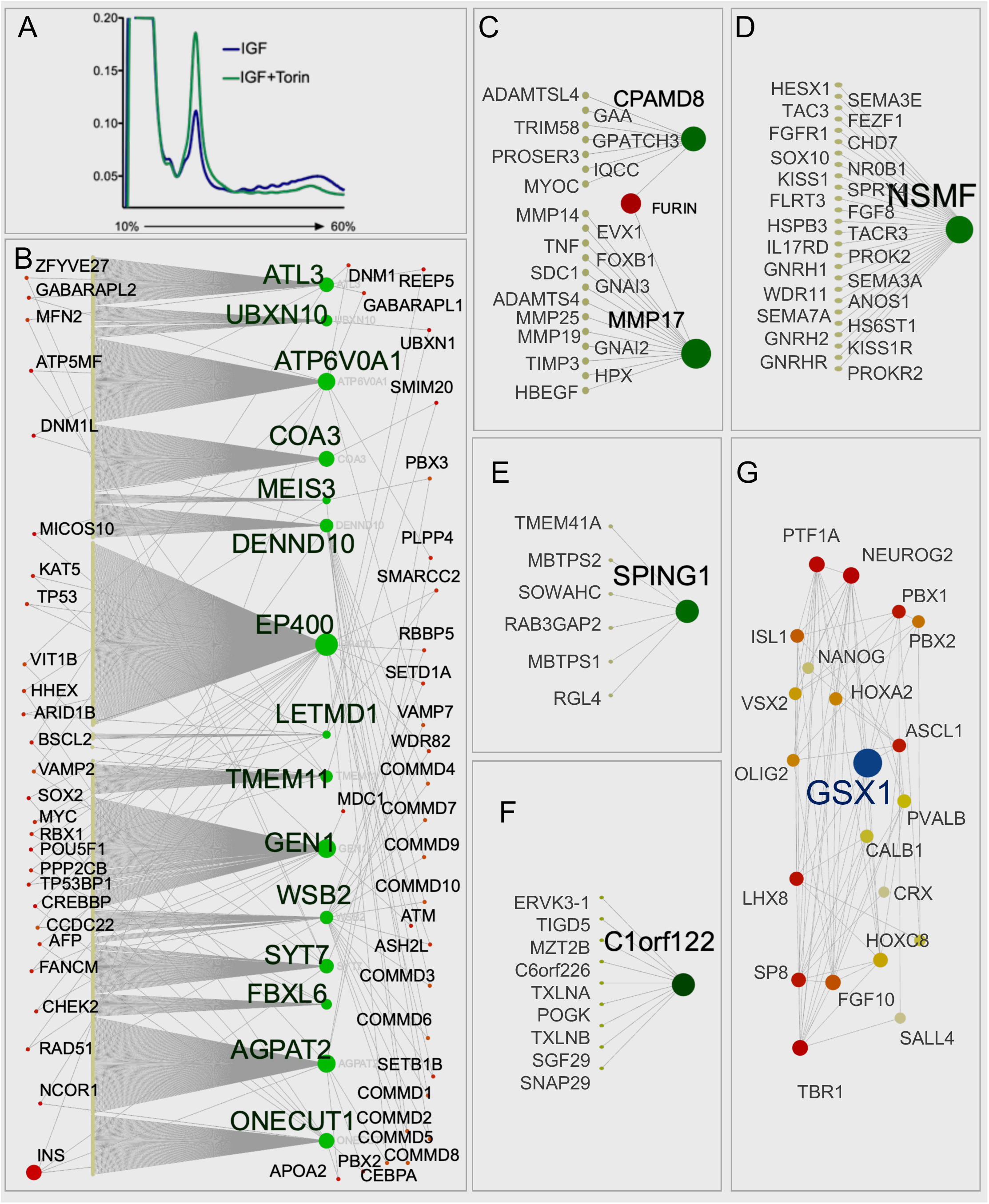
IGF1–mTOR signaling enhances translational control and reshapes protein interaction networks governing neurodevelopment, proteostasis, and cell-environment dynamics. **(A)** Polysome profiles show that IGF1 stimulation promotes polysome formation in FOXG1⁺ neural progenitors, whereas mTOR inhibition (Torin1) collapses polysome structures, indicating impaired translational capacity. **(B)** Protein-protein interaction (PPI) network built from IGF1-responsive translational targets highlights hub proteins (e.g., ATL3, UBXN10, EP400, SYT7) involved in mitochondrial function, vesicle trafficking, transcriptional regulation, and neurodevelopment. **(C)** Focused PPI network centered on CPAMD8 and MMP17 reveals enrichment for extracellular matrix remodeling and proteolytic regulation pathways important for neural progenitor niche dynamics. **(D)** NSMF-centered PPI network identifies connections to axon guidance, neuroendocrine development, and hypothalamic migration programs, suggesting IGF1 regulation of neuroendocrine axis formation. **(E)** SPING1-centered PPI network highlights interactions with ER stress response regulators (e.g., MBTPS1/2) and vesicle trafficking components, linking IGF1 signaling to proteostasis and membrane homeostasis. **(F)** uncovers interactors related to vesicle transport (TXLNA, SNAP29), cytoskeletal organization, and transcriptional regulation, suggesting coordinated control of intracellular trafficking and gene expression by IGF1-responsive translation. **(G)** GSX1-centered PPI network highlights interactions with ASCL1, LHX6, LHX8, NEUROG2, OLIG2, PBX1, PBX2, PTF1A, SP8, and TBR1, linking IGF1 signaling to forebrain neural differentiation.

## Materials and Methods

### Human Pluripotent Stem Cell Culture

Human pluripotent stem cell line H1 (WiCell) and iPSC (APOE3-BC1) and the APOE3-BC1 induced pluripotent stem cell (iPSC) line, were maintained on mitotically inactivated mouse embryonic fibroblast (MEF) feeder layers in standard human PSC medium. The original APOE4/4 BC1 iPSC line was generated by Dr. Linzhao Cheng (Johns Hopkins University) from human bone marrow–derived CD34⁺ cells via transient expression of a non-integrating plasmid^34^. Subsequent gene correction to the APOE3/3 genotype was performed in our laboratory. The medium comprised DMEM/F12 (Gibco, Cat# 11330-032), supplemented with 20% KnockOut Serum Replacement (Gibco, Cat# 10828-028), 4 ng/mL recombinant human FGF2 (PeproTech, Cat# 100-18B), 1 mM GlutaMAX (Gibco, Cat# 35050-061), 100 μM non-essential amino acids (Gibco, Cat# 11140-050), and 100 μM 2-mercaptoethanol (Sigma-Aldrich, Cat# M3148). Medium was replaced daily, and cells were passaged every 4–6 days using 1 mg/mL collagenase type I (Gibco, Cat# 17100-017) in DMEM/F12 at a 1:6 to 1:12 ratio. All procedures involving human ESCs were conducted in accordance with guidelines set by the Johns Hopkins University Institutional Stem Cell Research Oversight Committee.

### Derivation and Culture of Human FOXG1⁺ Forebrain Neural Progenitor Cells

To initiate neural differentiation, human ESC or iPSC colonies were pre-treated with 1 mg/mL collagenase type I (Gibco, Cat# 17100-017) in DMEM/F12 for 5–10 minutes at 37°C. As colony borders began to lift while centers remained attached, collagenase was removed and replaced with growth medium. Detached colonies, free of underlying MEFs, were carefully transferred to low-attachment six-well plates (Corning, Cat# 3471) and cultured in suspension for 2 days in human ESC medium lacking FGF2. From days 2 to 6, dual-SMAD inhibition was initiated by supplementing the medium (hereafter referred to as KoSR medium) with 50 ng/mL recombinant human Noggin (R&D Systems, Cat# 6057-NG-025), 1 μM dorsomorphin (Tocris, Cat# 3093), and 10 μM SB431542 (Tocris, Cat# 1614). On day 7, the resulting embryoid bodies (EBs) were transferred to plates coated with either Matrigel (Corning, Cat# 354277) or laminin (Sigma-Aldrich, Cat# L2020) and cultured in N2 induction medium composed of DMEM/F12 (Gibco, Cat# 11330-032), 1% N2 supplement (Gibco, Cat# 17502-048), 100 μM MEM non-essential amino acids (Gibco, Cat# 11140-050), 1 mM GlutaMAX (Gibco, Cat# 35050-061), and 2 μg/mL heparin (Sigma-Aldrich, Cat# H3149). Cultures were fed with N2 medium every other day from days 7 to 12, and daily thereafter. By days 8–9, attached EBs flattened and differentiated into neural rosettes. Continued induction led to the emergence of compact, columnar 3D neural aggregates—termed rosette-derived neural aggregates (RONAs)—in the centers of rosette colonies. These RONAs were manually microdissected with care to avoid contaminating peripheral flat cells or underlying layers, and maintained as neurospheres in Neurobasal medium (Gibco, Cat# 21103-049) supplemented with B27 minus vitamin A (Gibco, Cat# 12587-010) and 1 mM GlutaMAX for 24 hours. RONAs were then enzymatically dissociated into single cells and replated onto laminin/poly-D-lysine–coated plates (Sigma-Aldrich, Cat# P0899) for downstream assays.

### Matrigel Insert Cultures and Niche Modeling

To model niche-dependent IGF1 signaling, forebrain NPC aggregates were transferred onto Matrigel-coated permeable Transwell inserts (0.4 μm pore size, Corning, Cat# 3470) in six-well plates and maintained for up to 10 days with medium changes every 48 hours. Recombinant human IGF1 (PeproTech, Cat# 100-11) at 100 ng/mL, IGF1-neutralizing antibody (R&D Systems, Cat# AF-291-NA), or IGF1R antibody (Abcam, Cat# ab9572) were added to designated cultures to assess ligand dependency.

### IGF1 Manipulation and Proliferation Assays

For IGF1 manipulation, recombinant human IGF1 (PeproTech, Cat# 100-11) was added at 100 ng/mL. Neutralizing antibodies against IGF1 (R&D Systems, Cat# AF-291-NA) and IGF1R (Abcam, Cat# ab9572) were used at 5 μg/mL. For proliferation assays, aggregates were incubated with 10 μM BrdU (Sigma-Aldrich, Cat# B5002) for 2 hours prior to fixation. BrdU incorporation was visualized via immunostaining, and Ki67⁺ cell fractions were quantified in cross-sections. Caspase-3 immunostaining was used to assess apoptosis

### Immunofluorescence Staining and Imaging

NPCs or organoid sections were fixed in 4% paraformaldehyde (Electron Microscopy Sciences, Cat# 15710), permeabilized with 0.3% Triton X-100 (Sigma-Aldrich, Cat# T8787), and blocked with 5% donkey serum (Sigma-Aldrich, Cat# D9663). Primary antibodies included anti-FOXG1 (Abcam, Cat# ab18259), anti-Nestin (Millipore, Cat# MAB5326), anti-Ki67 (Cell Signaling Technology, Cat# 9129), anti-Caspase-3 (Cell Signaling Technology, Cat# 9661), anti-IGF1R (R&D Systems, Cat# AF305), and anti-CK8 (Developmental Studies Hybridoma Bank, Cat# TROMA-I).Fluorescent secondary antibodies (Alexa Fluor conjugates, Thermo Fisher Scientific, various catalog numbers) were applied, followed by DAPI nuclear counterstain (Thermo Fisher Scientific, Cat# D1306). Images were acquired using a Zeiss 880 Confocal Microscope System.

### Manders’ Overlap Coefficient (MOC) Quantifies the Degree of Spatial Colocalization

To quantify the degree of spatial colocalization between two markers, we calculated the Manders’ Overlap Coefficient (MOC). For instance, Nestin (green channel) and IGF1R (red channel), which evaluates the proportion of pixels exhibiting co-occurrence of fluorescence signals, independent of intensity correlation. Grayscale images representing Nestin and IGF1R immunofluorescence were first loaded and converted to floating-point format using the img_as_float function from the skimage Python library to ensure numerical precision. To enhance contrast and mitigate background variability, we applied adaptive histogram equalization (equalize_adapthist) to each image. Signal-positive regions were then identified through intensity thresholding using Otsu’s method, and binary masks were generated for each channel, where pixels above the threshold were assigned a value of 1 and background pixels a value of 0. The resulting binary masks for Nestin and IGF1R defined the signal distributions for subsequent overlap analysis.

To isolate colocalized regions, we generated a combined mask by performing a logical AND operation between the two binary masks, identifying pixels with co-detected Nestin and IGF1R signal. Manders’ coefficients were calculated from this combined mask. Specifically, M1 represents the fraction of Nestin-positive pixels that also contained IGF1R signal, and M2 represents the fraction of IGF1R-positive pixels overlapping with Nestin. These were computed using the following equations: M1 = ∑(Nestin ∩ IGF1R) / ∑(Nestin), and M2 = ∑(Nestin ∩ IGF1R) / ∑(IGF1R), where the sums reflect the number of signal-positive pixels. All steps were implemented in Python using the skimage and numpy libraries, enabling reproducible, threshold-aware quantification of spatial colocalization between the two fluorescent markers.

### Quantitative PCR (qPCR)

Total RNA was extracted using the RNeasy Mini Kit (Qiagen, Cat# 74104), and cDNA was synthesized with SuperScript IV VILO Master Mix (Thermo Fisher Scientific, Cat# 11756050). qPCR was performed with Power SYBR Green Master Mix (Thermo Fisher Scientific, Cat# 4367659) on a QuantStudio 6 Flex system (Applied Biosystems, Cat# 4485691). Primer sequences targeted FOXG1, PAX6, HES1, GSX1, and housekeeping controls (GAPDH, ACTB). Relative expression was calculated using the ΔΔCt method.

### Neurosphere Assay

Single-cell suspensions of NPCs were seeded at low density (1,000 cells/well) in ultra-low-attachment six-well plates (Corning, Cat# 3471) in neural expansion medium. IGF1, IGF1R inhibitors, or antibodies were added as specified. After 7 days, spheres were quantified by number and categorized by diameter (<100 μm, 100–200 μm, or >200 μm) using phase contrast microscopy and ImageJ analysis.

### RNA sequencing

Human FOXG1⁺ NPCs were harvested and total RNA was collected from culture following the manufacturer’s protocol. Libraries were generated using the Illumina®-compatible ScriptSeq™ mRNA-Seq Library Preparation Kit according to manufacturer’s instruction. The quality of the libraries was checked on the Bioanalyzer using a High Sensitivity DNA Chip (Agilent, Waldbronn, Germany) and quantified using the Illumina Genome Analyzer DNA library quantification kit (Kapa Biosystems, Woburn, MA). Quality-control tests for the unmapped reads were performed using the FASTQC software. The resulting reads were mapped to the GRCh38 annotated human reference genome using default settings. The aligned reads that were mapped to genes and defined by annotation were then counted with htseq-count. Reads were annotated using a custom R script using the UCSC known Genes table as a reference. Gene-to-GO term association data was obtained from the Gene Ontology of GO Consortium website (http://geneontology.org/). Genes assigned to a particular GO term were also assigned to their parent terms upward in the GO graph structure. GO term enrichment was evaluated using hypergeometric tests, and a p-value was estimated using a hypergeometric distribution.

Enrichment terms and associated genes and statistics are presenting with additional statistical analyses also provided. g:Profiler was applied to show functional enrichment analysis on input gene list, mapping known functional information sources and detecting statistically significantly enriched biological processes and pathways and highlighted in Manhattan plot [PMID: 31066453] (https://biit.cs.ut.ee/gprofiler). Gene ontology shows output from an enrichment analysis, searching the following categories: GO Biological Process, GO Molecular Function, PANTHER Biological Process, PANTHER Molecular Function, KEGG and PANTHER pathway. A web-based data platform is built for an interactive search of the preconditioning expression data measured under the NMDA and OGD preconditioning.

### Ribosome Profiling (Ribo-seq)

Human FOXG1+ NPCs were cultured under growth factor/insulin-deprived conditions for 16 hours before treatment with 20 ng/mL recombinant human IGF1 (PeproTech, #100-11) for 30 minutes, or pre-treated with 250 nM Torin 1 (Tocris, #4247) for 30 minutes followed by 30 minutes of IGF1 in the continued presence of Torin 1. This protocol allows detection of acute, mTOR-dependent translational responses. Cycloheximide (100 μg/mL; Sigma-Aldrich, #C4859) was added to halt translation elongation. Cells were lysed in polysome lysis buffer, and clarified lysates were treated with RNase I (Ambion, #AM2295) to generate ribosome-protected fragments. Monosomes were isolated via ultracentrifugation through a sucrose cushion at 100,000 x g for 4 hours using an SW41Ti rotor (Beckman Coulter). RNA from the ribosome pellet was extracted using Trizol LS (Invitrogen, #10296-028) and separated on a 15% TBE-urea polyacrylamide gel (Invitrogen, #EC6885BOX). Fragments between 25–32 nt were excised and purified. Library construction followed Ingolia et al. protocol with modifications and was validated using a Bioanalyzer 2100 (Agilent). Libraries were sequenced on an Illumina NovaSeq 6000. Reads were aligned using TopHat2 to the human genome (GRCh38), and gene-level FPKM values were computed from BAM files. Translation efficiency was defined as ribosome footprint FPKM / RNA-seq FPKM. Transcripts with ≥1.5-fold reduced TE in Torin-treated samples (n=2) were defined as IGF1-mTOR targets.

### Polysome Profiling and 5′UTR Reporter Assays

Cells were treated with IGF1 or Torin+IGF1, lysed in polysome buffer (20 mM Tris-HCl pH 7.5, 140 mM NaCl, 5 mM MgCl2, 1 mM DTT, 100 μg/mL cycloheximide), and loaded onto 10–60% sucrose gradients. Gradients were centrifuged at 35,000 rpm for 2 h at 4°C in an SW41Ti rotor (Beckman Coulter). Gradient profiles were recorded at 254 nm (Biocomp Gradient Station).

Fractions were collected and RNA was extracted using the miRNeasy Micro Kit (Qiagen, #217084). cDNA synthesis and qPCR were performed as above.

To assess 5’UTR-dependent translation, reporter constructs encoding Renilla luciferase under the control of GSX1, ACTB, or eEF2 5’UTRs were synthesized and cloned into pGL4 vectors (Promega). HEK293 cells were transfected with 5’UTR-luciferase constructs alongside a firefly luciferase normalization vector using Lipofectamine 3000 (Invitrogen, #L3000008). After 24 h, cells were treated with Torin 1 (250 nM) or DMSO for 6 h. Dual luciferase activity was measured using the Dual-Luciferase Reporter Assay System (Promega, #E1910). Relative Renilla activity was normalized to firefly and vehicle control.

### Dharmacon™ siRNA solutions and lentiviral transduction

GIPZ Lentiviral™ shRNA technology (Dharmacon™, Thermo Fisher Scientific, USA) was used to modulate gene expression in human neurons. Pre-designed human GIPZ shRNA Target Gene Sets comprised of 6 GIPZ shRNA constructs. Knockdown was compared to non-silencing control and experimental workflow was controlled using a pGIPZ non-silencing shRNAmir lentiviral vector (as a negative control). Lentivirus packaging was performed. GIPZ Human Lentiviral shRNA Clone Gene Set constructs targeting human gene GSX1 were cloned into lentiviral vector. Lentiviral vectors were co-transfected into HEK293FT cells with the lentivirus packaging plasmids pVSVg and psPAX2 using FuGENE® HD. Supernatants containing virus were collected 48– and 72-hours post transfection, passed through nitrocellulose filter (0.45 μm) and applied on cells in culture. At 12 weeks post-differentiation, human cortical neurons were transduced with lentivirus carrying control shRNA or targeting shRNA comprising complementary sequence of gene mRNA, was introduced into cells according to the manufacturers’ instructions (MOI of 5). Polybrene (5 µg/ml; Sigma-Aldrich) was added to the transduction medium to increase efficiency of the process. Cells transduced with non-silencing lentiviral shRNA vector (Thermo Fisher Scientific), containing a shRNA sequence with no homology to known mammalian genes, served as a negative control.

### Protein-protein interactions (PPI) network construction

The STRING database has been applied to search, analyze, and construct the protein-protein interactions (PPI) network (Search Tool for the Retrieval of Interacting Genes http://string-db.org/, Franceschini, A. et al. STRING v9.1: protein-protein interaction networks, with increased coverage and integration. Nucleic acids research). Full STRING network that indicates the edges of both functional and physical protein associations was used. The line color of network edges indicates the type of interaction evidence. The active interaction sources including text-mining, experiments, databases, co-expression, neighborhood, gene fusion, and co-occurrence were selected. The number of first shell interactors to show was set as no more than 5 interators. The number of nodes, number of edges, average node degree, average local clustering coefficient, expected number of edges, PPI enrichment p-value, and functional enrichments in the network (which include Biological Process, Molecular Function, Cellular Component, PubMed Reference publications, KEGG Pathways, Reactome Pathways, Tissue expression, Subcellular localization, Annotated Keywords, Protein Domains Pfam, Protein Domains and Features InterPro and SMART Protein Domains) were indicated in the **Table**. The combined score >0.4 was used as the cut-off criterion and was defined as significant protein– protein association networks with increased coverage, supporting functional discovery in genome-wide experimental datasets.

### Western Blot Analysis

FOXG1+ NPCs were lysed in RIPA buffer (Thermo Fisher, #89900) with protease and phosphatase inhibitor cocktail (Roche, #04693132001). Protein was quantified using the Pierce BCA assay kit (Thermo Fisher, #23225). Equal amounts (20–30 μg) were separated by SDS-PAGE (4–12% Bis-Tris gels, Invitrogen, #NP0321BOX), transferred to PVDF membranes (Millipore, #IPVH00010), and probed with antibodies against phospho-AKT (Ser473), total AKT, phospho-ERK1/2 (Thr202/Tyr204), total ERK1/2, phospho-S6 (Ser240/244), total S6, phospho-4EBP1 (Thr37/46), total 4EBP1, FOXG1, Pax6, HES1, and β-actin (all from Cell Signaling Technology). HRP-conjugated secondary antibodies (Jackson ImmunoResearch) and ECL reagent (Thermo Fisher, #32106) were used for detection. Bands were imaged using a ChemiDoc MP System (Bio-Rad) and analyzed with Image Lab software.

### Statistical Analysis

Statistical analyses were performed using GraphPad Prism (v9.5.1) and R (v4.3.1). All data are presented as mean ± standard error of the mean (SEM) unless otherwise stated. For comparisons between two groups, unpaired two-tailed Student’s t-tests were used if data met assumptions of normality and equal variance (confirmed using Shapiro–Wilk and F tests, respectively). For comparisons involving more than two groups, one-way or two-way ANOVA was used followed by Tukey’s post hoc test or Bonferroni correction when appropriate. For BrdU, Ki67, and neurosphere quantification, at least three independent biological replicates were used. Each replicate consisted of ≥10 aggregates or wells per condition.

Neurosphere sizes were binned into <100 µm, 100–200 µm, and >200 µm diameter, and analyzed by chi-square test or Kruskal–Wallis test if non-parametric.

qPCR data were normalized to GAPDH or ACTB as internal controls and analyzed using the ΔΔCt method. For ribosome profiling and RNA-seq, differential expression and translational efficiency were calculated using custom R scripts and Bioconductor packages (DESeq2, edgeR, Xtail), with Benjamini-Hochberg FDR correction applied to control for multiple testing. Manders’ overlap coefficients (M1 and M2) were calculated from binary colocalization masks using the skimage and numpy packages in Python. Differences in MOC values between conditions were assessed using two-tailed unpaired t-tests. Luciferase reporter assay data were expressed as Renilla/Firefly ratio normalized to vehicle control. At least three technical replicates were performed per condition, and results were analyzed by one-way ANOVA with Dunnett’s test. A p-value < 0.05 was considered statistically significant and denoted in figures as *p < 0.05, **p < 0.01, ***p < 0.001, \*\*\**p < 0.0001*, with exact p-values provided in figure legends.

## Notes

### Competing Interest Statement

The authors have declared no competing interest.

### Summary of Updates

This work was supported in part by the National Institute on Aging, National Institutes of Health, P30 AG066507 (Johns Hopkins ADRC). The content is solely the responsibility of the authors and does not necessarily represent the official views of the NIH.

### Funding

- National Institute on Aging: P30 AG066507

